# Investigating the effects of contextual information, visual working memory, and inhibitory control in hybrid visual search

**DOI:** 10.1101/2023.09.29.560143

**Authors:** Alessandra Barbosa, Anthony J. Ries, Juan E. Kamienkowski, Matias J. Ison

## Abstract

In real-life scenarios, individuals frequently engage in tasks that involve searching for one of various items stored in memory. This combined process of visual search and memory search is known as hybrid search. To date, most hybrid search studies have been restricted to average observers looking for previously well-memorized targets in blank backgrounds. Here we investigated the effects of context and the role of memory in hybrid search by modifying the task’s memorization phase to occur in all-new single trials. Additionally, we aimed to assess how individual differences in visual working memory capacity and inhibitory control influence performance during hybrid search. In an online experiment, 110 participants searched for potential targets in images with and without context. A change detection and go/no-go task were also performed to measure working memory capacity and inhibitory control, respectively. We show that, in target present trials, the main hallmarks of hybrid search remain present, with a linear relationship between reaction time and visual set size, and a logarithmic relationship between reaction time and memory set size. Context affected search efficiency in different ways. In target-absent trials we found large differences between context-present and absent conditions, suggesting participants’ adoption of adaptive strategies. Finally, working memory capacity did not predict most search performance measures. Inhibitory control, when relationships were significant, could account for only a small portion of the variability in the data. This study provides insights into the effects of context and individual differences on search efficiency and termination.

## Introduction

### Visual Search and Memory Search

Visual search (VS) is the action of looking for a target amongst distractors. It is a ubiquitous task for instance, when searching for products in stores. Mainly used to examine visual attention (Treisman & Gelade, 1980), VS was intensively researched for decades. The core manipulation in VS - varying distractor set sizes, where one target-object is nested, while measuring the time until target detection (reaction times or RTs), and measuring detection accuracy (Neisser, 1964). From these, a hallmark was established - the linear dependence between RT and visual set size (VSS; Wolfe, 2020).

Therefore, while much is known about elements that influence visual search when *one* sole item is searched for, search in real-life is substantially more complex, often involving several objects in memory. From Schneider and Shiffrin (1977) classic work, encompassing both visual search (VS) and memory search (MS), hybrid search (HS) is when observers search for any of many possible targets (Wolfe, 2012b). HS tasks involve memorizing potential targets to subsequently identify their incidence in a display. In Schneider and Shiffrin (1977), different memory manipulations included whether HS had consistent-mapping (when target sets are fixed, used throughout trials), all-new mapping (when new items compose the memory set each trial), or varied-mapping (when targets become distractors and vice-versa), which negatively impacted efficiency and accuracy in this order.

Regardless of the specific conditions, a robust logarithmic relationship between RT and memory set size (MSS) has been consistently reported: up to MSSs of 100 (Wolfe, 2012b), for categories (i.e., even larger MSSs; Cunningham & Wolfe, 2014), and for words (Boettcher & Wolfe, 2015). Nonetheless, most recent HS studies have not manipulated memory-mappings. Solely utilizing consistent-mapping-like paradigms, high accuracies are observed (e.g., see Wolfe, 2012b). Under such conditions, speed–accuracy trade-offs that vary across set sizes cannot be evaluated as adequately as by paradigms that limit memory strengths (Nosofsky, Cox, et al., 2014). Given the prominent role of context in-memory representations, such as recognizing a face or place after a single real-world encounter (Ison et al., 2015), examining paradigms with trial-by-trial changing memory sets is justified.

### Search Models

To explain the behavioral signatures observed, models for HS drew from VS and MS models. Cunningham and Wolfe (2014) postulated a 3-stage model for HS. After the first stage of *guided* search (Wolfe, 2007), where solely feature-plausible objects affiliates to the memory set are considered for the second stage of object recognition, a third stage of logarithmic MS is performed through a diffusion process. Each memorized target item races as information is accumulated in parallel over time, forming N-diffusion processes, where N=MSS. Target selection only occurs when information from a target accumulator reaches the decision threshold. The threshold is set higher to impede exaggerated false alarms and lower to promote speed - the speed-accuracy trade-off (Chun & Wolfe, 1996).

Target absent conditions, when search must be terminated (i.e., search ends without target localization), still needs better clarification (Drew et al., 2017). In VS, The Competitive Guided Search (CGS) model of Moran et al. (2013) provides a robust basis to understand search termination, explaining it as a quit unit added in the information accrual phase. This quit unit *competes* with the remaining visual set and is continuously adjusted depending on the likelihood that the target is yet to be found in the current stimuli. If the quit unit is selected, search is terminated. This adaptation would be compatible with HS studies’ results for VS (Wolfe, 2020); and can aid understanding of HS termination processes.

Additionally, as most HS paradigms follow a condition akin to consistent-mapping, not much is known when targets’ memory strengths are manipulated (Wolfe et al., 2015). Drawing from MS literature, which focuses on the processes underlying MS through memory-mapping manipulations, Nosofsky, Cox, et al. (2014) proposed an exemplar-familiarity random-walk MS model with a logarithmic RT diffusion stage. In contrast to recent HS models, it proposes that racing items are dictated by their ‘memory strength’, influenced by repetition effects and time presented, while they compete, and information is accumulated. Thus, while the curvilinear relationship of RTxMSS is maintained, efficiency and accuracy are affected (Nosofsky, Cao, et al., 2014; Nosofsky, Cox, et al., 2014), which can elucidate HS processes for different memorization conditions.

### Context in Hybrid Search

Moreover, most HS findings are based on experiments conducted with artificial stimuli on blank backgrounds (Wolfe, Võ, et al., 2011), which may limit their ecological validity. In VS, accumulating evidence shows that context is critical in guiding attention in real-world sear (Wolfe, Alvarez, et al., 2011). Indeed, context guidance was found to overpower bottom-up saliency in guiding eye movements in naturalistic search (Henderson et al., 2009); and to facilitate search in scenes for targets in plausible locations rather than implausible ones (Neider & Zelinsky, 2006; Wolfe et al., 2011) or blank backgrounds (Wolfe, Alvarez, et al., 2011). However, naturalistic scenes are complex/continuous, with VSSs impossible to define (Rosenholtz et al., 2007) but they are never random (Henderson & Ferreira, 2004). Scenes have syntax, i.e., structural plausibility - humans appear on horizontal superficies (Biederman, 1976; Torralba et al., 2006); and semantics, i.e., meaningful associations - for example, a toothbrush on a sink (Võ & Henderson, 2009).

In HS, the limited exploration of contextual information has left mixed results. To our knowledge, only Boettcher et al. (2013) and Boettcher et al. (2018) attempted to examine contextual effects in HS. However, both were investigating whether context aided memory set partition to context-relevant items at *fixed* VSSs and MSSs, rather than understanding its possible impacts on established RT signatures and efficiency as VSSs and MSSs increase. Whereas Boettcher et al. (2013) found evidence suggesting memory search could be partitioned by context trial-by-trial (i.e., constraining the memory search to context-relevant-only items); Boettcher et al. (2018) reported that overall individuals seemed to be incapable, or reluctant, to restrict memory sets trial-by-trial, but were able to partition search to memory subsets that continued relevant for multiple trials.

### Visual Working Memory and Search

Visual working memory (VWM) entails the maintenance and manipulation of a limited amount of visual information that serves current task demands (Luck & Vogel, 1997). Resource-limited, there is converging behavioral evidence estimating capacity limits around 3-4 items (Cowan, 2001; Vogel & Machizawa, 2004) with individual differences in capacity being significantly predictive of higher cognitive function measures (Luck & Vogel, 2013). VWM has been largely implicated in a variety of VS processes, such as representing the search template to guide attention (Desimone & Duncan, 1995; Soto et al., 2006), comparing the target template to potential suitor objects (Bundesen, 1990), and categorizing stimuli (Duncan & Humphreys, 1989). Given this, individual differences in VWM capacity should be expected to play a significant role in HS.

Nevertheless, individual differences in VWM capacity have rarely been assessed directly in HS. Even in research examining the average searcher, equivocal results are observed. Dual-task studies in VS and HS, where search is combined with a change detection task (CDT), a measure of VWM, have shown mixed results. Studies in VS have found that loading VWM with CDTs did not affect VS performance in several conditions (Oh & Kim, 2004; Treviño et al., 2021; Woodman et al., 2001), except when *spatial* visual memory was filled (Downing & Dodds, 2004; Oh & Kim, 2004; Woodman & Luck, 2004), or when targets changed in each trial (Woodman et al., 2007). Congruently, in HS, Drew et al. (2016) found that loading VWM did not affect HS efficiency, albeit HS did diminish VWM *capacity* by a fixed amount regardless of VSS and MSS variation.

### Inhibitory Control and Search

Inhibitory control (IC) is the critical executive function of suppressing goal-irrelevant stimuli interference and of subduing prepotent motor response (Young et al., 2018). Experimental paradigms conventionally include brusque prepotent response incitation, where one either proceeds or subdues action (e.g., go-no-go tasks; Miyake et al., 2000). Individuals who have higher false alarms, or larger negative response bias, when subjected to signal detection theory (SDT) analysis, have more difficulty inhibiting prepotent responses (Young et al., 2018).

Since Treisman and Sato (1990) proposed a *feature-based inhibition*, VS models demonstrated the importance of IC in top-down selection by filtering-out goal-irrelevant distractors with ‘templates for rejection’, avoiding irrelevant attentional capture, and in preventing return to searched locations (inhibition of return) (Baithwaite et al., 2005; Beck & Hollingworth, 2015; Gaspelin & Luck, 2018). In target selection, Usher and McClelland (2001) also propose lateral inhibition is at play between target evidence accumulators, where increased inhibition leads to increased accuracy and RT spread. In target absent conditions, Moran et al. (2013) model also assumes the operation of inhibitory links between the priority map and the quitting unit to select search termination. Thus, inhibition’s role in VS performance is argued critical.

Taken together, it is surprising that individual differences in inhibitory control are overlooked in HS literature (Clarke et al., 2020). Sparse evidence exists in foraging search. While Jóhannesson et al. (2017) and Ólafsdóttir et al. (2019) have found no relationship between individual differences in IC and foraging patterns/ability (e.g., foraging speed and target-template switching), in both studies IC was measured through a complex inhibition task, confounding working memory in the measurement (see Jóhannesson et al., 2017; Ólafsdóttir et al., 2019).

### Present Study

Whilst the well-known linear relationship of RT and VSSs, and the logarithmic dependency between RT and MSSs, have been replicated under different conditions (Boettcher & Wolfe, 2015; Wolfe, 2012b), a gap in the literature remains regarding the potential effects of contextual information in HS. Moreover, most recent HS research has emulated a consistent-mapping paradigm, which has been seen to produce near error-free data (Nosofsky, Cox, et al., 2014). Promoting a trial-by-trial change of target items, akin to an all-new mapping manipulation, may help better observe speed-accuracy trade-offs. This manipulation would also be critical to further uncover memory’s role in search, since VWM involvement might only be observed when targets change per trial (Woodman et al., 2007), and it is still unclear in MS whether context can/cannot restrict the memory set to scene-relevant items on a trial-by-trial basis (Boettcher et al., 2013, 2018). The investigation of individual differences in VWM capacity would complement such examination by directly measuring it, extending its previous scarce understanding in HS (Drew et al., 2016). Indeed, literature still concentrates on how the average searcher performs (Clarke et al., 2020). Given the proposed importance of IC in VS models for top-down selection and search termination (Moran et al., 2013; Treisman & Sato, 1990), an evaluation of the potential impact of individuals’ IC is also merited. Therefore, along with investigating the effects of context and brief memorization in HS’ behavioral markers, this study aims to examine the potential effects of VWM capacity and IC.

Based on previous literature, some predictions with regards to the study manipulations can be made. With mounting replications of HS’ RT signatures in divergent conditions, all-new memorization and context are not expected to *qualitatively* affect HS’ RT signatures but affect efficiency and accuracy. Memory strength of targets are seen to be weaker in all-new conditions in comparison to consistent-mapping manipulations’ fixed, repeated target set (Donkin & Nosofsky, 2012). Whereas lower accuracies in general might be observed, context present conditions might provide higher accuracies and lower RTs in comparison to context absent conditions, given context has been seen to guide search (Torralba et al., 2006), and, in some conditions, facilitate MS by partitioning search to context-relevant items (Boettcher et al., 2018).

Further, whereas VWM interacts with VS (Desimone & Duncan, 1995), its actual role in HS is more ambiguous. If this study follows Drew et al. (2016) account that a fixed-amount of VWM is used as a conduit to transfer incoming target templates to memory, one might expect VWM capacity to correlate with RT intercepts in HS, where individuals with higher VWM capacity might transfer targets through VWM faster than lower VWM-capacity individuals. However, given our modified paradigm with targets changing per trial, analogous to Woodman et al. (2007), this proposition might not hold and we might see higher VWM capacity producing smaller RT slopes as well; given high-VWM capacity individuals would have higher storage capacity/resource allocation flexibility, as set sizes increase (Luck & Vogel, 2013). Concerning IC, given its potential importance in many VS processes and search termination (Moran et al., 2013), in HS, individuals with higher IC are expected to have larger RT intercepts than lower IC individuals, reflecting a potential to better maintain conservative thresholds with less FAs and higher accuracy.

## Methods

### Participants

An online data collection method gathered data from 110 participants, identifying as: women (59); men (49); non-binary (1); different identity (0); and non-disclosing (1). Participants were recruited via email and social media. Their ages (excluding 3 age misreports) ranged from 18 to 61 years-old (*M=*27 and *SD=*7.63). Data were also collected for another 10 participants that were excluded due to aborting the experiment (N=7) or low behavioral performance (N=3). Participants were given the option to enter a draw for Amazon vouchers worth £20 each so two people were compensated. Convenience sampling was adopted, as email and social-media outreach is limited to users and not randomized. Online recruitment concentrated on Chinese-oriented social media platforms (WeChat) and in Brazilian social media groups. The study was approved by the University of Nottingham School of Psychology Ethics Panel (ethics approval: S1240).

### Materials

The experiments were implemented in Psychopy and executed online by participants through Pavlovia.org (Peirce et al., 2019). Albeit having some variability in personal screen sizes, the display was set to 1920 x 1080 pixels resolution, only running in laptops or desktop computers. Previously tested and considered the most stable, Google Chrome (MacAskill et al., 2022) was requested of participants before the experiment. MATLAB (MATLAB, 2020) and IBM SPSS (version 26.0; IBM-Corp, 2019) were used for data analysis.

Information and consent slides were shown pre-experiment for individuals to read and consent participation by pressing “y”. After experiment completion, de-briefing slides appeared. In Part 1 of the experiment, participants searched 112 images of 1, 2, 4, or 8 stimuli with (N=56) or without (N=56) context after memorizing a set of 1, 2, 4, or 8 stimuli. All targets and distractors belonged to the same category and items only appeared once during the experiment. Target and distractor images were composed of either animals, objects, or people, whereas contextual imagery encompassed open spaces (e.g., forests) and closed spaces (e.g., shelves). Across the 112 images shown, 62 included animals ((subcategories: dog, cat, bear, horse, cow, sheep, bird, wild animals, farm animals), 38 included objects (subcategories: toys, cup, vase, bottle, kite, book, remote, teddy bear, plant, fruits) and 12 included people, therefore categories were not equally represented. Target/distractor images were resized so that their new size would be compatible with the background image and placed according to scene syntax (i.e. no major violations of support, interposition, position, and size). Locations on context absent trials were the same as in context present trials. All of these were photorealistic stimuli built from images from COCO dataset (Tsung-Yi et al., 2014) and ImageNet (Deng et al., 2009), being then digitally combined with background images.

Part 2 of the experiment encompassed assessing individuals’ VWM capacity and IC. VWM capacity was measured using the classic Change Detection Task (CDT) adapted from Balaban et al. (2019) and Xu et al. (2018). It involved 120 trials with set sizes 4 (N=60) and set size 6 (N=60) of colored blocks. Changes occurred in 50% of the trials. To measure IC, the well-established go/no-go (GNG) task was employed. It was based on the parameters of Young et al. (2018) and Wessel (2018). To isolate IC and tackle GNGs’ issue of confounding other cognitive processes, the 150 trials had solely 1 stimulus - a blue circle in go trials (N=120) and an orange circle in no-go trials (N=30). Thus, a 4:1 ratio of go/no-go trials was maintained.

### Procedure

Random participant IDs were either assigned inside recruitment emails or generated during the experiment if no ID was inputted. The larger experiment was split in 2 parts (or sessions) for partakers’ convenience. Part 1 involved the HS and part 2 encompassed the CDT and GNG task. Due to practical ID assignment issues, as well as to maintain peak participant concentration in the main task (session 1), counterbalancing of tasks between sessions was not done.

### Hybrid Search Task

When searchers clicked on the dedicated URL-link for part 1, a pop-up window pre-experiment required participants’ ID and age. Afterwards, the experiment progressed to the study’s information sheet, consent form, and instructions in black text and white background (Arial, 0.1cm letter height, position [0,0]). When consenting and understanding to press “m” for target present and “v” for target absent, participants pushed the “s” key to start the HS. The trial begins with 1, 2, 4, or 8 to-be-memorized images, the MSS, on a display. A fixation cross follows (position [0,0]) until the search screen is shown with 1, 2, 4, or 8 images, the VSS, to be searched from (see Figure 1). They contained at most 1 memory set item. Subsequent trials promptly initiated after a response, or, after 7 seconds if no response is given. Target and context were present in 50% of trials. Completion of the 112 trials automatically triggered the de-briefing slides. Trials alternated between one of the 4 combinations of conditions - *either target present or target absent conditions in a context absent or context present trial*. Completion time was around 20 minutes.

**Figure 1.**
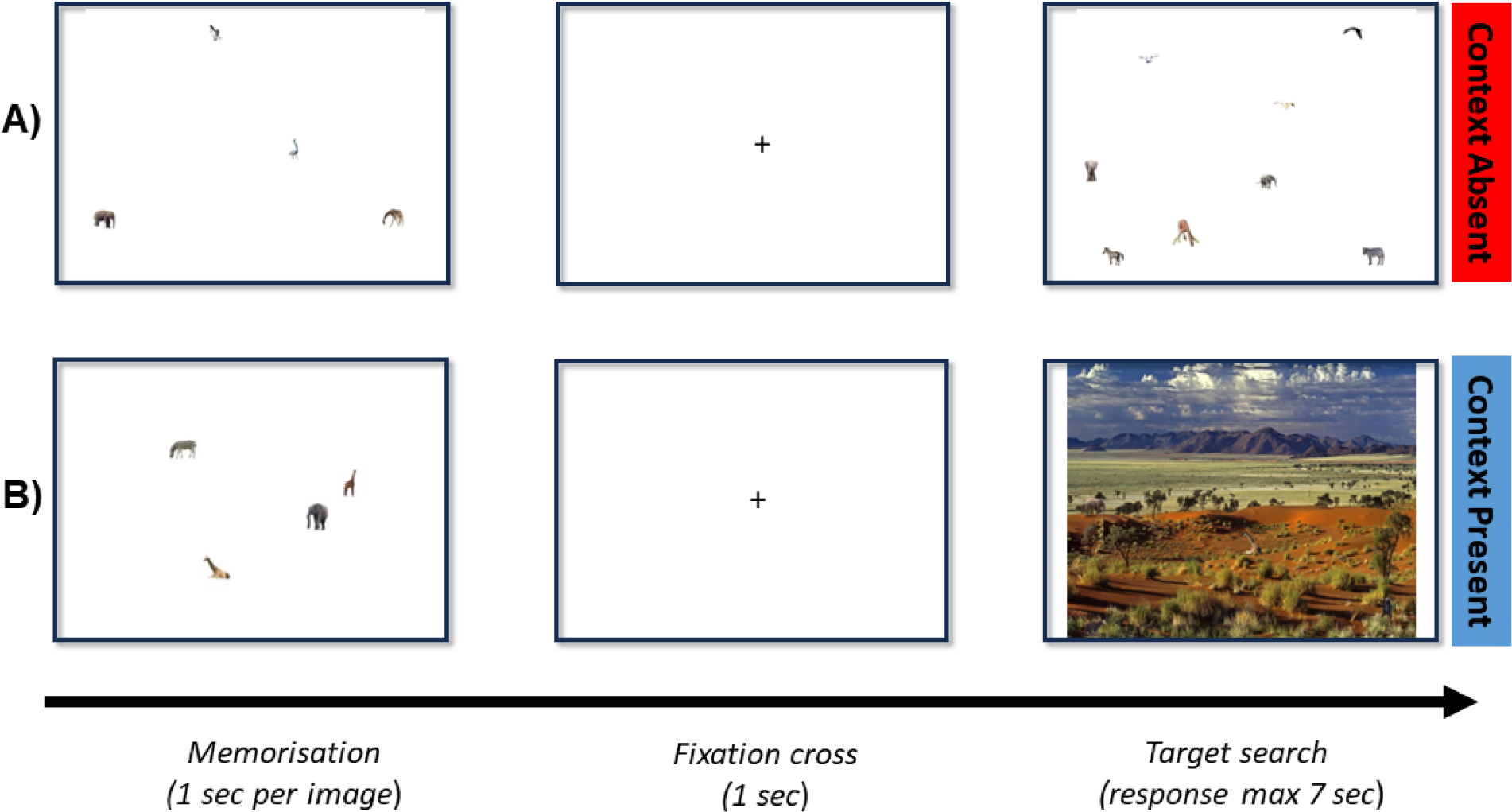
*A) Model trial for a memory set size of 4 and a visual set size of 4 without contextual information. B) Model trial for a memory set size of 4 and a visual set size of 4 with context.* Each trial initiates with the images to be memorized, which is then followed by a fixation cross. The search screen (containing or not the target) follows until a response is made.

### Change Detection Task (CDT) and Go-No Go Task (GNG)

Participants were allowed to break and return for the experiment’s second part, englobing the CDT and GNG task. Clicking the link, participants input their assigned participant ID. Subsequently, information and consent sheets were shown. The CDT’s instructions (Arial, 0.03cm letter height, position [0,0]) appeared explaining the task and informing to foveate on the fixation cross. There were 4 training trials pre-experiment without feedback, which included a message in-between trial refreshing the instructions. Trials began with 4 or 6 color-blocks, followed by fixation cross and, subsequently, a detection screen with 1 color-block (for timeframes see the *Supplementary Information, figure S1*). If the same color and location as in the initial display appeared, “k” key was pressed. If a different color and/or location popped up, “l” was pushed. The next trial automatically started after a response without feedback. Changes occurred in 50% of trials. Once the 120 CDT trials were completed (approximately 10 minutes), the GNG task instructions appeared. After 4 practice trials with feedback, the GNG task started by displaying either blue (go-stimuli; size [0.15, 0.15]cm) or orange circles (no-go-stimuli; size [0.15, 0.15]cm) at a 4:1 go/no-go ratio. For go-stimuli the space bar was pressed and for no-go-stimuli that action had to be inhibited. With an intertrial interval of 450ms, 150 trials lasted approximately 3 minutes (see the *Supplementary Information, figure S2*). The experiment terminated with de-briefing and voucher draw instructions. Part 2 lasted approximately 15 minutes in total.

### Design & Data Analysis

For counterbalancing, trial order was randomized across participants in each experimental task. Each participant experienced a unique sequence. The data was retrieved from Pavlovia.org to Excel files and matched by participant ID. Participants that aborted within session, in-between sessions, and had outlier performances that departed more than 3 *SDs* from the mean across conditions were discarded.

First, a descriptive analysis was conducted to assess the data’s statistical metrics to provide validation of the experiments’ properties and distributions delivered in an online medium. Second, a preliminary analysis was conducted to satisfy data assumptions necessary for the main analyses. Pre-analysis checks examined outliers, homoscedasticity, normality of residual errors, and sphericity by inspecting Cook’s distance, the distribution of the residual error, and Mauchly’s test. Where Mauchly’s test indicated that the sphericity assumption had been violated (here, for variables with 2+ levels), Greenhouse–Geisser correction was applied. When significant and variance-covariance matrices were not homogenous, conservative Pillai’s Trace was reported. No outliers (maximum Cook’s distance surpassing 1) were identified. For simple linear regressions, collinearity was assessed. No correlations above r>.9 were observed. By visually inspecting scatterplots of residuals against independent variables, no systemic pattern was observed (homoscedastic). Also, examining residuals’ histograms, all conformed to an acceptable departure from normality.

Third, the main analyses that address this study’s aims were undertaken. To attend to our first objective, linear and logarithmic regression fits were constructed in MATLAB for VS and MS RT/Accuracy as a function of set size. ANOVAs further quantified the relationships. Consistent with HS literature, this study considers the F-test robust enough against Type I errors in non-normal data (Blanca Mena et al., 2017). The relationship between accuracy and RT was directly examined through linear regression for VS and MS. Wilcoxon Signed Rank Tests were also employed to determine the logarithmic fit superiority for MS, as well as search efficiency differences in context present conditions versus context absent, as these directly compared Goodness of Fits, fitted intercepts, and fitted slopes.

Moreover, to address our second objective, simple linear regressions were performed to assess the potential effects of individual differences in VWM capacity and IC on HS performance. Here, the independent variables include K and c; and dependent variables involve: MS’ and VS’ RT slopes, RT intercepts, accuracy, and FAs. Age was also checked for its potential impact in search; however, it did not show a significant result.

While some measures in this study are from direct observations, VWM capacity and IC are measured by K and c, respectively (see Supplementary Information). K is the estimate of an individual’s VWM capacity, whereas c is seen as a measure of decision/response bias. These are calculated through Cowan’s (2001) formula, and through SDT analysis (Green & Swets, 1966), respectively.

## Results

### Online Validation of Experiments

Descriptive statistics of hybrid search, CDT, and GNG tasks show properties consistent to previous lab-based experiments in literature, conferring adequate online validation. Congruous to Wolfe (2012b), this task saw a mean accuracy well above chance level (M = 0.70, SD = 0.06). The characteristic decline in accuracy when VSSs/MSSs increase in divergent memory-mapping conditions was also observed: VSS1=0.8 to VSS2=0.56; and MSS1=0.93 to MSS8=0.54. Thus, in good agreement with recent results of context-lacking HS tasks (Drew et al., 2017).

Visual working memory capacity K was calculated as in Balaban et al. (2019) (see Supplementary Information Figure S1 for the task sketch and main formulas). K was considered normally distributed after a K-S normality test was performed and was not significant D (110) =.065, p>.05. To be meaningful, K is expected to depict the underlying normally distributed VWM capacities in population (Balaban et al., 2019). Further, this study’s K replicates 2 pivotal characteristics concerning VWM capacity – (1) capacity-limited characteristic (M = 2.31); and (2) significant individual differences denoted by *SD* = 0.82 (Balaban et al., 2019). It is consistent with China-based (K:2.14; Xu et al., 2018) and American studies (K:2.55; Fukuda et al., 2016). These findings support the validity of our online CDT results’.

SDT analysis was applied to calculate GNG’s decision bias measure, denoted as c. A negative c mean (M = -0.23, SD = 0.58) was achieved, consistent with existent literature (Young et al., 2018).

### Context Effects and Trial-by-trial Memorization in Hybrid Search

In this section, the following abbreviations will be used for orderliness - target present (TP); target absent (TA); context present (CP); context absent (CA).

For RT analyses, from a total of 12,320 trials across 110 participants, we excluded trials (N=217) with no answer within the maximum time allowed (7 seconds). Very short responses (less than 200 ms) were only recorded for 4 trials and were not excluded from the data. To quantify RT search patterns, two 3-way 4×2×2 repeated-measures ANOVA were performed to further assess the relationship between either VSSs/MSSs (1 x 2 x 4 x 8), target presence (present x absent) and context presence (present x absent) on RT. The same was done for accuracy as the dependent variable. Partial eta square (ηp^2^) was disclosed as effect size throughout. Following (Cohen, 1988), ηp^2^ >.06 is considered medium, and ηp^2^ >.14 a large effect size. Further, to capture potential speed-accuracy trade-offs, a linear regression was performed to quantify the relationship between accuracy and RTs.

Figure 2 exhibits the regression fits constructed for RT as a function of VSSs/MSSs. The classic linear dependence between RT and VSS was evident (TP/CA R^2^=.70; TA/CA R^2^=.73; TP/CP R^2^=.53; TA/CP =.40). Also, the positive logarithmic relationship between RT and MSS was observed (TP/CA R^2^=.66; TA/CA R^2^=.52; TP/CP R^2^=.66), except in TA/CP condition, which saw a negative logarithmic function (TA/CP R^2^=.40). Correct-trial RTs, shown in Fig. S3, saw main effects of VSS [F (2.63,186.74)= 140.3, p<.001, *ηp^2^=.*66], target presence [F (1,71) =116.39, p<.001, *ηp^2^=.*62], and context presence [F (1,71) =108.32, p<.001, *ηp^2^=.*60]. These interacted significantly, F (2.64,187.35) =17.17, p<.001, *ηp^2^=.*20. Main effects’ pairwise comparisons showed RT means were significantly different from each other in every VSS (p<.001). Also, RTs in TA were significantly larger than TP, and in CA were significantly smaller than CP conditions (all p<.001). These were Bonferroni-corrected for multiple comparisons (at p<.05). Simple main effects (SME) analysis was employed after significant interaction effects and is available inside *Supplementary Information, table S1*. Briefly, all VSSs, in TA and TP conditions, CA had smaller RT means than CP conditions, apart from VSS4 TA and VSS8 TA conditions, where CA and CP RTs did not significantly differ.

**Figure 2.**
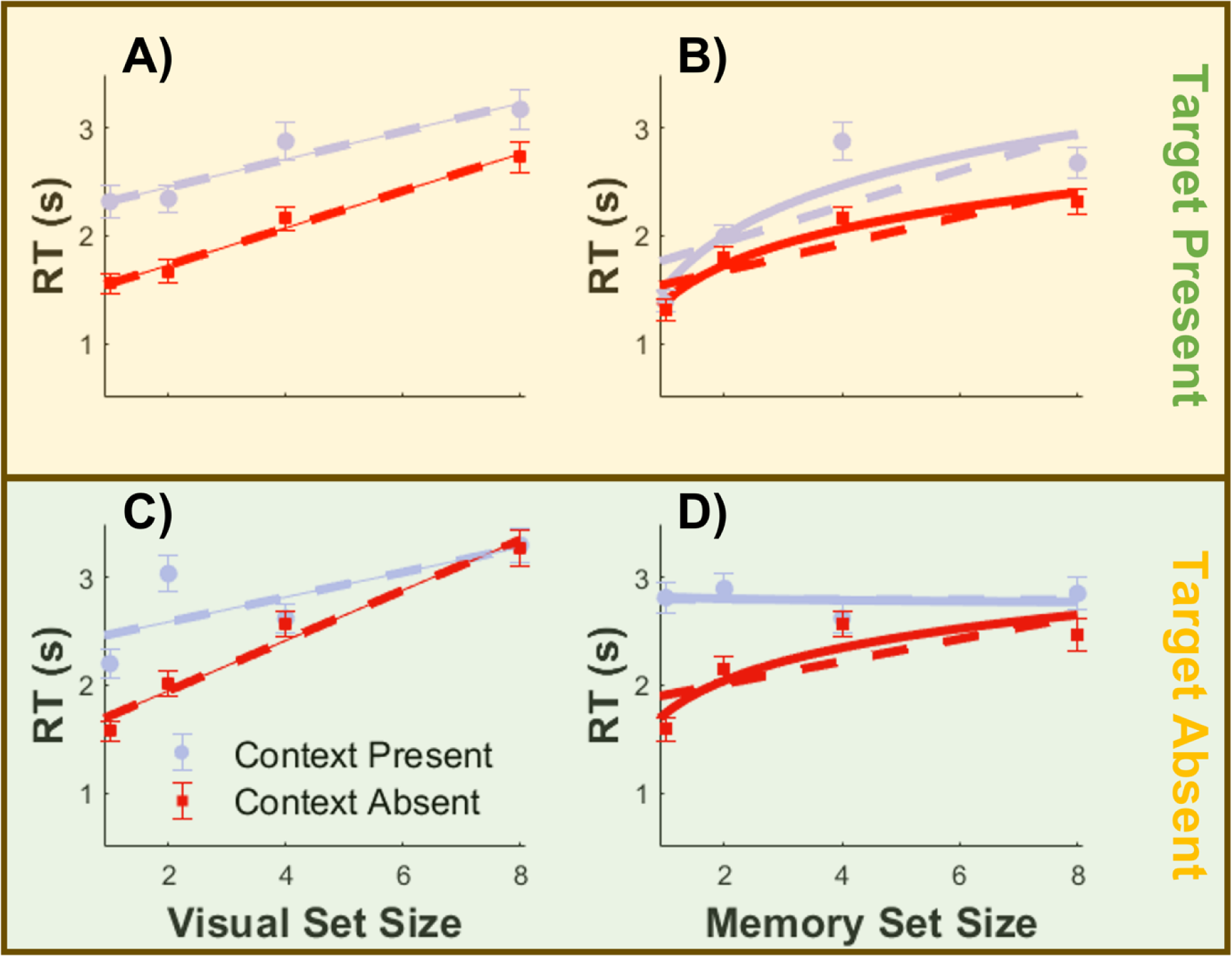
*RT as a function of visual and memory set sizes for TP, TA, CP and CA conditions.* *Note.* Error bars denote 95% CI. Dashed lines depict linear fits, while continuous lines portray logarithmic fits. Panels differentiate conditions: A) Visual Search Target Present; B) Memory Search Target Present; C) Visual Search Target Present; D) Memory Search Target Absent. Equations for each condition – VS: y = 0.17*x + 1.38 (TP/CA); y = 0.12*x + 2.19 (TP/CP); y = 0.23*x + 1.48 (TA/CA); y = 0.12*x + 2.36 (TA/CP). MS: y = 0.34*log(x) + 1.39 (TP/CA); y = 0.47*log(x) + 1.52 (TP/CP); y = 0.31*log(x) + 1.74 (TA/CA); y = -0.015*log(x) + 2.82 (TA/CP). In RT x VSS, MSS was fixed at SS4; in RT x MSS, VSS was fixed at SS4.

Similarly, MSS [F (2.49,184.27) =89.85, p<.001, *ηp^2^=.*55], target presence [F (1,74) =162.89, p<.001, *ηp^2^=.*69], and context presence [F (1,74) =129.32, p<.001, *ηp^2^=.*64] had main effects on correct-trials RT. These interacted significantly, F (2.25,166.5) =35.84, p<.001, *ηp^2^=.*33. Bonferroni-corrected for multiple comparisons (at p<.05), main effects’ pairwise comparisons for MSS, target presence, and context presence, on correct-trial RTs was performed. All RTs were significantly larger as SS increased (all p<.001), except for MSS4 and MSS8 (p>.05). Also, TA had larger RT means than TP, and CA had faster RTs than CP condition (all p<.001). Following significant interaction effects, SME analysis was employed and is displayed in *Supplementary Information, table S2*. At all MSSs TA and TP conditions, CA had significantly faster RTs than CP conditions, except at MSS4 TA condition, where CA’s and CP’s RT were not significantly different.

Across all conditions, targets were missed on 34% of target-present trials, and false-alarms were produced on 27% of target-absent trials. The corresponding discriminability d’ was 1.02. There were main effects of accuracy VSS [F (2.54,276.99) =123.95, p<.001, *ηp^2^=.*53], target presence [F (1,109) =16.13, p<.001, *ηp^2^=.*13], and context presence [F (1,109) =3.36, p=.069, *ηp^2^=.*03]. Interactions were significant, F (2.71,295.38) =11.14, p<.001, *ηp^2^=.*09. Corrected with Bonferroni, main effects’ pairwise comparisons showed that accuracy significantly diminished as VSSs rose, except at VSS2 to VSS4, where it significantly increased (p<.001). TA had significantly higher accuracy (p<.001) than TP conditions, whereas CA and CP did not significantly differ on accuracy (p=.069). Following significant interactions, SME analysis was employed. See *Supplementary Information, table S3*. At VSS2 TP, VSS8 TA and TP, CA had significantly greater accuracies than CP conditions. Contrastingly, at VSS2 TA, VSS4 TA and TP, CA had significantly lower accuracies than CP.

In MS, there were main effects of MSS [F (2.35,256.48) =438.29, p<.001, *ηp^2^=.*80], target presence [F (1,109) =23.09, p<.001, *ηp^2^=.*18], and context presence [F (1,109) =8.36, p=.005, *ηp^2^=.*07] on accuracy as well. They interacted significantly, F (2.53,275.97) =10.53, p<.001, *ηp^2^=.*09. Bonferroni-corrected main effects’ pairwise comparisons of MSS, target and context presence, were performed. As MSSs rose, accuracy significantly declined (all p<.001). TA had significantly higher accuracy than TP conditions, and CA saw significantly lower accuracies than CP conditions (all p<.005; see Figure 3). SME was employed after significant interaction effects, see *Supplementary Information, table S4,* for details. When interactions significantly differed, CA conditions had lower accuracies than CP conditions, except at MSS4 TP condition.

**Figure 3.**
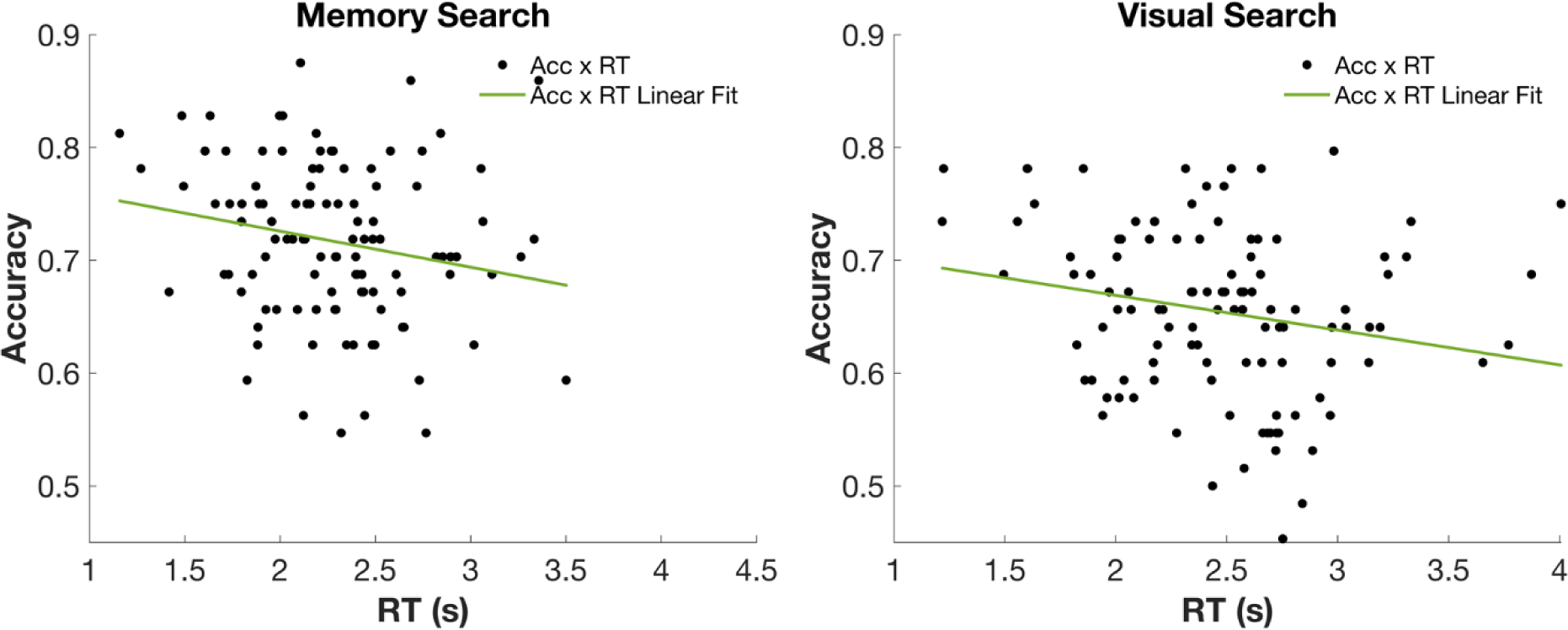
*Response accuracy as a function of reaction times.* Left panel: Accuracy x RT in Visual Search (MSS locked at set size 4); Right panel: Accuracy x RT in Memory Search (VSS locked at set size 4).

To further illustrate the relationship between accuracy and RT, a linear regression was performed for both VS, F(1, 108) = 5.107, p = .026, R^2^ = .045, R^2^ adjusted = .036, and MS, F(1, 108) = 4.508, p = .036, R^2^ = .040, R^2^ adjusted = .031. The relationships were significant such that, in VS, an increase of one second in RT meant that accuracy decreased by about 3.1% (B = -.031, 95% CI[-.033,-.028]); and, in MS, that accuracy decreased by about 3.2% (B = -.032, 95% CI[-.035,-.029]). Figure 3 displays accuracy as a function of RT for VS and MS.

### Memory Search: Log or Linear?

Moreover, to validate whether the RTxMSS relationship indeed follows a linear or logarithmic function, Wilcoxon signed-rank tests were performed. Figure 2 and table 1 show logarithmic fits significantly explained more RT variance than linear fits in all MS’ conditions (all p<.001), except TA/CP condition (p>.05).

**Table 1.** Wilcoxon Signed Rank test results comparing linear and logarithmic goodness of fits for memory search functions.

| Conditions | Goodness of Fit (Mean R2) |  |
| --- | --- | --- |
|  | Linear | Log |
| TP/CA | .55** | .66** |
| TP/CP | .50** | .66** |
| TA/CA | .43** | .66** |
| TA/CP | .41 | .40 |

*Note.* Wilcoxon Signed Rank Test – $p \leq .05 = *$ ; $p \leq .001 = **$ . Target Present – TP; Target Absent – TA; Context Present – CP; Context Absent – CA.

### Search Efficiency: Context Absent (CA) and Context Present (CP)

To investigate potential effects of context presence on search efficiency, Wilcoxon rank-signed tests were performed on RT slopes and intercepts. Table 2 shows that whilst VS ‘and MS’ RT intercepts when context was present were significantly larger than intercepts when context was absent, irrespective of target presence, RT slopes when context was absent were significantly less efficient than in context-present conditions, apart from the RT slope during memory search when target was present (all p<.05).

**Table 2.** *Search efficiency: comparing Context Absent (CA) and Context Present (CP) conditions*.

|  |  | TP |  | TA |  |
| --- | --- | --- | --- | --- | --- |
|  |  | CA | CP | CA | CP |
| **Visual Search** | Intercept | 1.38** | 2.19** | 1.48** | 2.36** |
|  | Slope | .17* | .12* | .23** | .12** |
| **Memory Search** | Intercept | 1.39** | 1.52** | 1.74** | 2.82** |
|  | Slope | .34** | .47** | .31** | -.015** |

*Note.* Wilcoxon Signed Rank test – $p \leq .05 = *$ ; $p \leq .001 = **$ ; Intercept and slopes of individual participant's fits; Target Present – TP; Target Absent – TA; Context Present – CP; Context Absent – CA

### Memory Search in Target Absent Condition: False Alarms

Given the unexpected RTxMSS pattern in memory search when the target was absent and the context present (TA/CP condition), an analysis of false alarms (FAs) is warranted to better understand search termination. A two-way 4×2 repeated-measures ANOVA was performed to investigate MSS and context presence effects on FAs. There were main effects of MSS [F(2.22,241.64)=340.43, p<.001, ηp^2^=.76], and context presence [F(1,109)=11.67, p=.001, ηp^2^=.10] on FA. They interacted significantly [F(2.4,256.17)=12.63, p<.001, ηp^2^=.10].

Bonferroni-adjusted (at p<.05), main effects’ pairwise comparisons of MSS and context presence on FAs were performed. MSS1 and MSS2 did not significantly differ (p>.05), while MSS4 had significantly less FAs than MSS8 (p<.001). CA had significantly more FAs than CP conditions (p=.001).

SME analysis, following significant interaction, is displayed in Table 3. At MSS1 [F(1,109)=3.85, p>.05, *ηp^2^=.*03] and MSS2 [F(1,109)=.01, p>.05, *ηp^2^=.*00], CA and CP did not significantly differ, while at MSS4 [F(1,109)=6.32, p<.05, *ηp^2^=.*06] and MSS8 [F(1,109)=20.35, p<.01, *ηp^2^=.*16], CA had significantly more FAs than CP conditions.

**Table 3.**
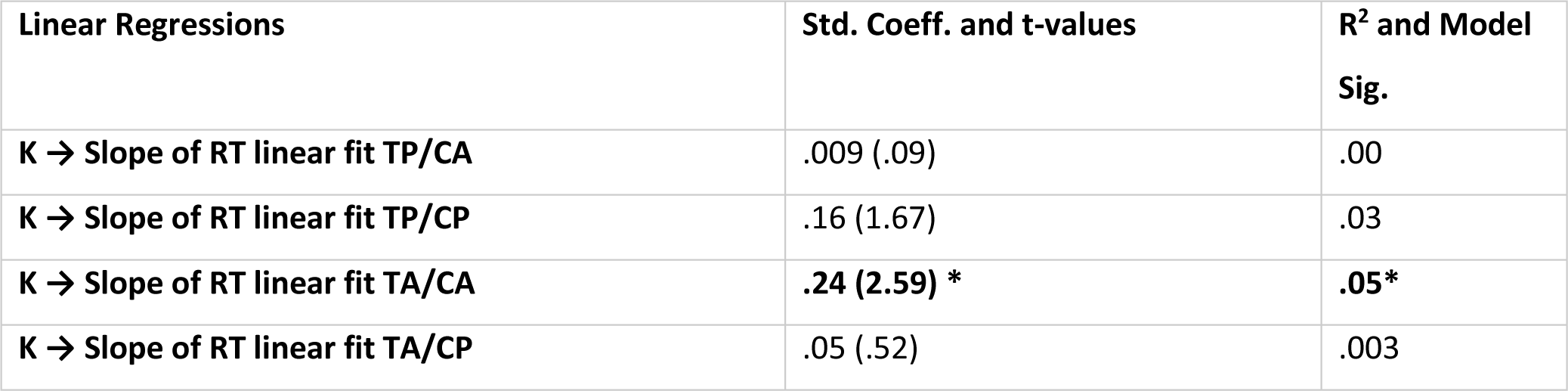
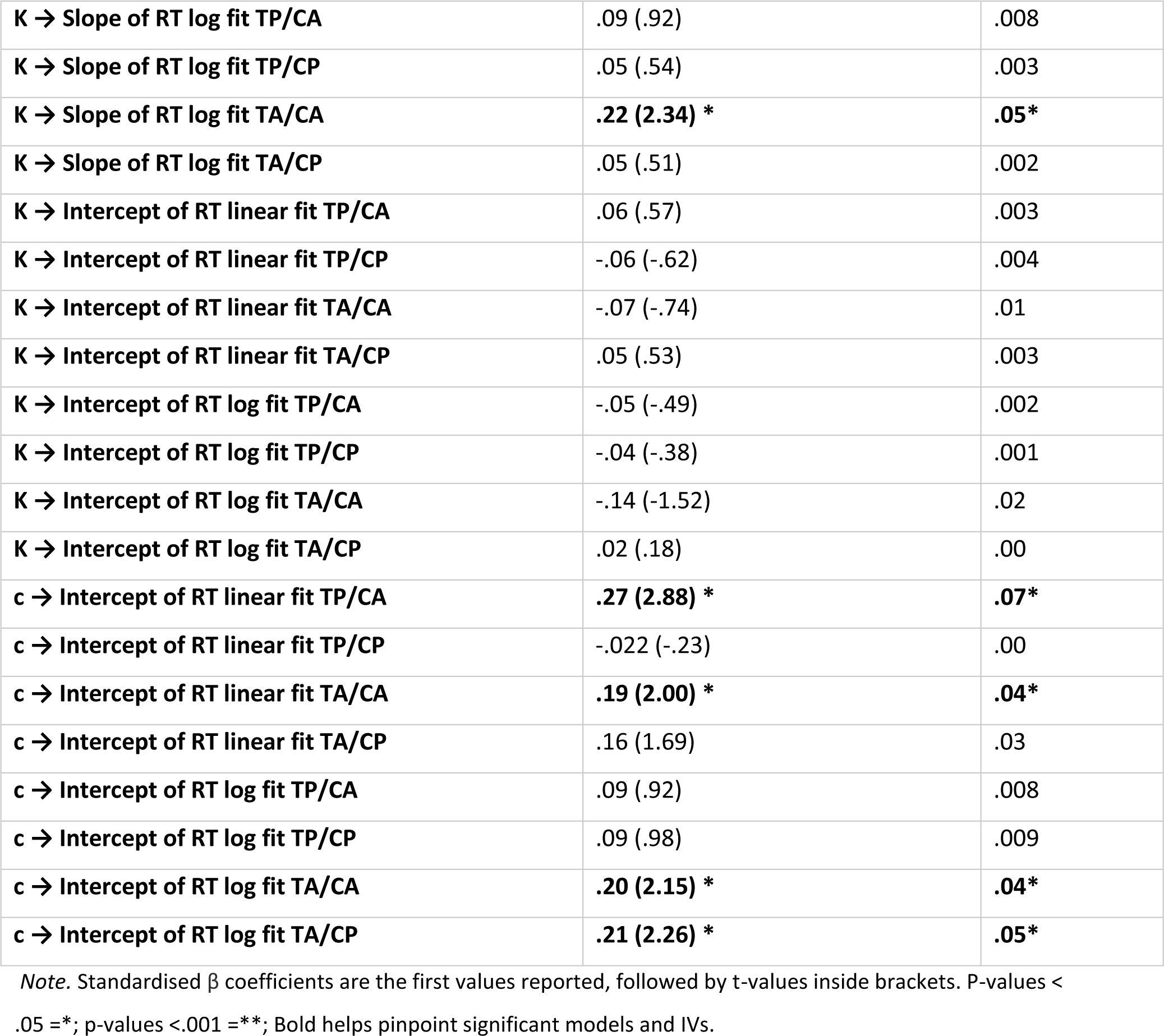
Simple Linear Regressions of K and c on RT data.

| Linear Regressions | Std. Coeff. and t-values | R<sup>2</sup> and Model Sig. |
| --- | --- | --- |
| K → Slope of RT linear fit TP/CA | .009 (.09) | .00 |
| K → Slope of RT linear fit TP/CP | .16 (1.67) | .03 |
| K → Slope of RT linear fit TA/CA | **.24 (2.59) \*** | **.05\*** |
| K → Slope of RT linear fit TA/CP | .05 (.52) | .003 |
| K → Slope of RT log fit TP/CA | .09 (.92) | .008 |
| K → Slope of RT log fit TP/CP | .05 (.54) | .003 |
| K → Slope of RT log fit TA/CA | **.22 (2.34) \*** | **.05\*** |
| K → Slope of RT log fit TA/CP | .05 (.51) | .002 |
| K → Intercept of RT linear fit TP/CA | .06 (.57) | .003 |
| K → Intercept of RT linear fit TP/CP | -.06 (-.62) | .004 |
| K → Intercept of RT linear fit TA/CA | -.07 (-.74) | .01 |
| K → Intercept of RT linear fit TA/CP | .05 (.53) | .003 |
| K → Intercept of RT log fit TP/CA | -.05 (-.49) | .002 |
| K → Intercept of RT log fit TP/CP | -.04 (-.38) | .001 |
| K → Intercept of RT log fit TA/CA | -.14 (-1.52) | .02 |
| K → Intercept of RT log fit TA/CP | .02 (.18) | .00 |
| c → Intercept of RT linear fit TP/CA | **.27 (2.88) \*** | **.07\*** |
| c → Intercept of RT linear fit TP/CP | -.022 (-.23) | .00 |
| c → Intercept of RT linear fit TA/CA | **.19 (2.00) \*** | **.04\*** |
| c → Intercept of RT linear fit TA/CP | .16 (1.69) | .03 |
| c → Intercept of RT log fit TP/CA | .09 (.92) | .008 |
| c → Intercept of RT log fit TP/CP | .09 (.98) | .009 |
| c → Intercept of RT log fit TA/CA | **.20 (2.15) \*** | **.04\*** |
| c → Intercept of RT log fit TA/CP | **.21 (2.26) \*** | **.05\*** |

Note. Standardised $\beta$ coefficients are the first values reported, followed by t-values inside brackets. P-values < .05 =\*; p-values <.001 =\*\*; Bold helps pinpoint significant models and IVs.

### Individual Differences in Hybrid Search

Simple linear regressions were performed to investigate the impact of individual differences, in K and c, on HS performance: RT intercepts and slopes, accuracy, and FAs. Table 3 summarizes the regression results for RT data. Please see *Supplementary Information, table S5,* for the simple linear regressions results of individual differences on accuracy and FAs.

K did not significantly predict accuracy in any condition, except from TP/CP, as well as false alarms. However, in VS’ and MS’ TA/CA conditions, a higher K predicted larger RT slopes. Moreover, c did not predict either accuracy or FAs. Participants’ c did significantly predict VS’ RT intercepts in every CA condition, i.e., when target was absent and present, as well as all of MS’ RT intercepts in TA conditions, i.e., when context was absent and present, except in MS’ target present condition. Nevertheless, all of these explained little variability in HS performance (max. R^2=^.10).

## Discussion

This study aimed to investigate the potential effects of context and trial-by-trial memorization on HS performance and assess the possible interplay of VWM capacity and IC. Results replicated the characteristic RT signatures seen in the existent lab-based literature (see (Wolfe, 2020)), as well as the typical decrease in accuracy as set sizes increased, consistent with results observed in previous HS studies (Drew et al., 2017). Generally, as RT rose, accuracy diminished. Trial-by-trial memorization and context did not *qualitatively* affect RT. Whereas in visual search RT increased linearly as VSS rose, in memory search RT increased logarithmically as MSS escalated, with log fits significantly explaining more variance than linear fits. This was true except for the memory search target absent condition, where large differences were seen when context was present – i.e., a “negative” logarithmic RTxMSS relationship. With less false alarms made in context present conditions in larger set sizes, this may suggest an adaptive change of search strategy.

Further, while individual differences in visual working memory capacity did not predict HS performance in most conditions, inhibitory control impacted search when context was absent in visual search and when target was absent in memory search. Although significant, differences in inhibitory control explained very little of the variance in performance. Individuals with lower inhibitory control seem to have a higher chance of precocious search termination, as observed by lower RTs.

### The Effects of Single-trial Memorization and Context in Hybrid Search

#### Target Present: Context Absent

In target present conditions, as expected, results *qualitatively* replicate the characteristic RT signatures of HS even when context was present, and memorization occurred trial-by-trial. In VS, the canonical linear relationship between RTxVSS seen in serial VS literature and HS literature was replicated (Treisman & Gelade, 1980; Wolfe, 2007, 2012b); whereas in MS, the logarithmic relationship between RTxMSS seen in traditional MS studies and hybrid search was also observed here (Boettcher & Wolfe, 2015; Nosofsky, Cox, et al., 2014; Wolfe, 2012b). As anticipated, memorization of all-new targets trial-by-trial does not seem to qualitatively affect the way attention is deployed and visual information is processed in non-efficient VS (Shiffrin & Schneider, 1977).

Indeed, Schneider and Shiffrin (1977) demonstrated how manipulating memory lists to change (all-new mapping) or remain the same (consistent-mapping conditions - as seen in Wolfe (2012b) and existent HS literature) on every trial provokes dramatic differences in accuracy patterns and RT efficiency in HS, but not RT *signatures*. A previously memorized target set used repeatedly throughout VS trials can create memory reinstatement effects in MS - i.e., helps increase distinction between memory representations and distractors (Nosofsky, 1986). Nosofsky, Cox, et al. (2014) found mean RTs are much faster and error rates are much lower in the consistent-mapping condition than in an all-new condition in an *MS* task. Consistently, this study observed, in general across conditions, lower accuracy in comparison to previous HS studies, which had pre-tested fixed memory sets throughout trials (e.g., Cunningham & Wolfe, 2014; Wolfe, 2012b). Congruent with Shiffrin and Schneider (1977) seminal work, when memory mappings *differ* from consistent-mapping conditions, the relationship between accuracy and VSS/MSS tend to negatively mirror the relationship observed between RTxVSS/RTxMSS, due to strength differences in memory representations. When encoded trial-by-trial, the ability to attend to each memory set item’s unique features is reduced as MSSs increases (Mewhort & Johns, 2000).

#### Target Present: Context Present

Context influenced the efficiency of hybrid search. When context was present, we observed higher RT intercepts and smaller RT slopes compared to context-absent conditions. This finding aligns with studies demonstrating that context, even when scenes are arbitrary, does provide facilitation as VSSs increase (Võ & Wolfe, 2015; Neider & Zelinsky, 2006; Torralba et al., 2006; Võ & Henderson, 2009; Wolfe, Alvarez, et al., 2011). The observed pattern is consistent with studies proposing a two-stage process in VS. The first stage involves global processing (Buetti et al., 2016), where context may slow down this initial stage but subsequently facilitates serial attention until target recognition (Wolfe, Võ, et al., 2011). Hence, in contrast to random display of isolated objects, scene context provides valuable global information that guides eye movements towards high-probability regions of the scene (Malcolm & Henderson, 2010). Indeed, by using eye-tracking, Drew et al. (2017) found evidence that visual search and recognition in HS likely occur through one visual search with multiple memory searches.

Whereas context in our stimuli did not violate scene syntax, it did not systematically follow scene semantics. This approach differs from previous studies, where objects placed in scenes with semantic violations were identified to hamper scene guidance when compared to semantic-abiding objects in scenes (Wolfe, 2020). Despite targets/distractors being scaled and superimposed over background images, this study still found context to improve VS efficiency when compared to blank backgrounds, suggesting the added benefit of syntactic and occasionally semantic-matching contextual information. This is congruent with Biederman et al. (1982) that has seen syntax play a greater role than semantics of scenes for object identification. However, future studies using stimuli set with varying degrees of semantic association between the context and memory set items are needed to uncover the influence of context on hybrid search.

Previously, it was proposed that context could facilitate MS by effectively reducing the functional MSS - restricting MS to a part of the memory set where context and item are relevant to each other (Boettcher et al., 2013). However, this was proposed based on experiments where the memory set was kept constant throughout trials. In contrast, when the memory set switched per trial, for both arbitrary contexts and semantically associated context to the memory set, search RTs were as effortful as if the whole set in memory was searched, congruent with our results (Boettcher et al., 2018).

While the results support a potential resistance to partition memory (Boettcher et al., 2018), there can be alternative explanations. For instance, differently than Boettcher’s experiments, targets were memorized in blank backgrounds, not in their associated background. Hollingworth (2005) found that when targets were contained in the previewed scene, accuracy was reliably higher than targets were memorized in blank displays. Similarly, Evans and Wolfe (2022), using old/new recognition tasks, reported that accuracy sensitivity declined when items were memorized in one scene and tested in another. Thus, object representations seem to be tightly bound by spatial context and item-specific positions. Therefore, here, the subsequent search for items in another blank array, rather than embedded in a new scene, might have been easier.

#### Target Absent

In target and context absent conditions, RT shapes replicated the linear dependence between VSS and RT, and the logarithmic dependence between MSS and RT (Boettcher & Wolfe, 2015; Wolfe, 2012a). Accuracy overall also decreased as VSSs/MSSs increased. This pattern of search termination is consistent with the proposed HS model by Cunningham and Wolfe (2014), where search proceeds until no items remain with activations that are above a set threshold. The remaining items are deemed unlikely to be targets, are not visited by serial attention and not searched through memory (Chun & Wolfe, 1996; Drew et al., 2017). This threshold is sought to be set adaptively – conservative to minimize errors and liberal to minimize RT – but is dependent on several factors such as the value of search and crowding in VS (Wolfe, 2012a), and memory strength and memory set items’ similarities/dissimilarities in MS (Bae & Luck, 2017; Nosofsky, Cao, et al., 2014).

Nonetheless, the presence of context dramatically changed these RT signatures in MS. A “negative” logarithmic shape was observed, possibly suggesting an adaptive change of search strategy. While false alarms rose as MSS increased, that was still significantly lower than false alarms in MS’ target and context absent trials that replicated the positive logarithmic RTxMSS relationship. Therefore, if applying a speed-accuracy trade-off lens (Reppert et al., 2018), it would seem that context presence provided some form of facilitation, since context present trials saw less false alarms in larger set sizes (4 and 8), while also having lower RT slopes than when context was absent. Whereas both had decisions made around the same time, ∼2.5 – 3 seconds, participants in context present trials identified more correctly the target’s absence. This is concordant to research that finds context to increase accuracy in object recognition (Palmer, 1975) and recall (Josephs et al., 2016), especially when related objects are in the scene (Davenport, 2007). Nonetheless, FAs did drastically rise as MSSs increased. Therefore, there might be alternative explanations (see Limitations). Interestingly, Botch et al. (2023) recently explored the relationship between visual search efficiency in a classic search task and a naturalistic task using virtual reality technology. They reported a significant relationship in the search efficiency between both tasks when targets were present, but not on target absent trials. In visual search, context was seen to facilitate search. In memory search, context did not seem to aid the search through memory, but, when targets were absent, it influenced search strategy. Throughout conditions, a negative relationship between accuracy and RT was observed, which demonstrates the significance of different memory-mapping manipulations in the speed-accuracy trade-off outcomes.

### The Effects of Individual Differences in Visual Working Memory capacity and Inhibitory Control in Hybrid Search

#### Visual Working Memory (VWM) and VWM capacity

The general lack of relationship between VWM capacity and HS performance measures is largely inconsistent with accounts that propose that search/target templates reside in VWM to (1) bias attentional deployment to goal-relevant objects (Desimone & Duncan, 1995; Soto et al., 2008), and (2) be compared to suitor objects (Bundesen, 1990). If that was so, higher VWM capacity individuals, supposedly having larger storing capacities or better ability in manipulating attentional resources (see Luck & Vogel, 2013), would search more efficiently as set sizes increase, than observers with lower VWM capacity (Sobel et al., 2007).

Nonetheless, these results are partially consistent with the sparse HS literature that previously assessed VWM capacity’s role. Drew et al. (2016), using a dual-task paradigm with CDTs in-between HS tasks, found that performing a HS task diminished VWM capacity by a fixed amount (e.g., one slot) regardless of VSS and MSS variation. Hence, if it is assumed that a “one-item” channel/path must successively move visual items to LTM, VWM interaction would not be dependent on set size. That is, only a fixed amount of each individual’s VWM capacity would be used for the task. This would be consistent with the largely non-significant results seen here between VWM capacity differences and HS’ RT *slopes*.

Furthermore, an important difference between this study and Drew et al. (2016) is that memory sets here were not kept constant, changing trial by trial. In VS literature, some dual-task studies indicated that loading VWM, with a CDT embedded in a VS task, solely impacted search when target sets changed per trial, not when they were kept constant (Woodman et al., 2007; Woodman & Arita, 2011). Behaviorally they simply observed a slowing of RT slopes when loading VWM, but electrophysiological data suggested an effect in VWM representation maintenance. While the results presented are inconsistent with these findings, this points to the importance of evaluating electrophysiological data alongside behavior to pull apart meaning of behavioral observations.

The lack of relationship between VWM capacity and search observed is not a rare finding. It accords with Kane et al. (2006) that, through several demanding experiments, found VWM capacity to be irrelevant to performance; with Irons and Leber (2016) that found the same in a dynamically-changing VS; and with Jóhannesson et al. (2017) that saw VWM capacity not correlate with RT and other foraging search patterns. Nonetheless, maybe it is the case that only certain types of working memory affect search. Oh and Kim (2004) and Woodman and Luck (2004) found evidence that loading spatial working memory slows scrutiny in VS, whilst loading feature working memory did not (Oh & Kim, 2004; Woodman et al., 2001). Our experiment measured VWM capacity, without distinguishing spatial or feature visual working memory capacity. Future studies could measure both and examine whether different types of VWM ability affect HS.

#### Inhibitory Control

Studies investigating the effect of individual differences in inhibitory control (IC) on search, especially those with direct IC measures (Clarke et al., 2020), are markedly scarce. Given inhibition’s clear importance in VS (Beck & Kastner, 2009) and search termination (Moran et al., 2013), this study investigated the IC’s role in HS. Results showed that, in VS, individuals with higher IC (lower negative bias) had significantly higher RT intercepts than participants with lower IC when target was absent and present, but context was absent. In MS, this relationship was seen solely in target absent conditions, regardless of context. Additionally, IC did not relate to accuracy or FAs.

With IC impacting mostly target absent conditions, these results suggest that more inhibitory control is needed to remain in search when there is no target on the visual display. This is consistent with Moran et al. (2013) CGS model that advances inhibitory links between the priority search map and the quitting unit that terminates search when selected, and other MS models that propose a diffusion stage with laterally-inhibiting racing target-items (i.e., Cunningham & Wolfe, 2014;

Nosofsky, Cox, et al., 2014). In target present trials, the activation of the priority map is automatically increased, which leads to higher inhibition of the quit unit in comparison to target absent trials (Moran et al., 2013). In other words, target absent conditions inherently increase quitting probability.

The role of IC was also relevant in target *present* trials, but only when context was *absent*. This may point to the increased need of inhibition when search is not facilitated by context providing cue/guidance (Mertes et al., 2016). In MS, where context did not facilitate performance in target present conditions, we only see IC affecting target absent conditions.

### Limitations

While this study has made contributions towards a better understanding of search in real life, it also had some limitations. First, this study did not systematically manipulate the categories in memory sets, as well as scene semantics. Object representations in short-term memory are seen to interact based on their similarities and depend on their assigned attentional priority by means of cues, such that the characteristics of the prioritized/cued item influences others’ representation (Bae & Luck, 2017). Unaccounted category differences in memory sets and scene semantics (which could have provided cues) may have played a role in our mixed findings of context’s effect in MS. Second, the experiment design does not allow direct comparisons between the efficiency and accuracy in all-new mapping and consistent mapping conditions as this would require a separate comparison group. Since the mapping of the stimuli was not manipulated, the comparisons made here are related to the qualitative nature of the relationship between reaction times and set sizes. Moreover, the amount of time observers fixated on *individual* items was not controlled. This can also affect memory strength and VWM consolidation for each target item (Donkin & Nosofsky, 2012). Future studies could include eye-tracking to better understand how memory strength impacts MS, via dwell times, and how observers encode target items in memory. This can help disentangle VWM’s role in HS and understand memory strengths impact, which is not controlled for in the existent HS corpus (Nosofsky, Cox, et al., 2014).

Furthermore, as an online study, environmental factors were uncontrolled and attentional engagement throughout the experiment could not be assessed. This could have impacted search in several ways including participants’ appraised value of continuing search if there were external attentional demands present. Contextual information in search should be further investigated in lab-based studies to strengthen results observed.

## CONCLUSIONS

This study contributed to the understanding of different elements involved in searching in typical real-world situations, when more than one possible target is being searched for, contextual information is present, and searching is conducted after a single encounter. Briefly, whereas the manipulation of context and trial-by-trial memorization produced a robust replication of Hybrid Search reaction times signatures - established through analysis of correct-only trials, it impacted search efficiency and accuracy in different ways. This was true, except for memory search, where context presence produced a different reaction times structure when targets were absent. While individual differences in visual working memory capacity were not seen to affect hybrid search performance, participants with higher inhibitory control seemed to be able to avoid precocious search termination. In conclusion, our study reveals the intricate interplay of various behavioral mechanisms in hybrid search. While the observed results provide valuable insights, some key questions remain unanswered, particularly concerning the influence of scene semantics on hybrid search, as well as the dynamic interactions between visual and memory search, working memory, and inhibitory control. To address these intriguing aspects, future research can leverage recent technical advancements, such as concurrent M-EEG and eye movements recordings (Care et al., 2023; Dimigen & Ehinger, 2021). The integration of these cutting-edge methodologies promises to unveil the underlying physiological mechanisms driving hybrid search processes, further deepening our understanding of this complex cognitive phenomenon.

## ACKNOWLEDGEMENTS

The authors would like to thank Mingqian Luo for contributing towards data collection. This research was supported by CONICET (PIP 11220150100787CO) and ARL (Cooperative Agreement Number W911NF1920240 and W911NF2120237 awarded to MJI and JEK).

## List of Abbreviations

VS: Visual search
MS: Memory search
HS: Hybrid search
VSS: Visual set size
MSS: Memory set size
VWM: Visual working memory
IC: Inhibitory control
RT: Response time *Only in the Results section:*
TA: Target absent
CA: Context absent
TP: Target present
CP: Context present
SME: Simple Main Effects

## Supplementary information

Supplementary materials include additional data analyses.

### Availability of data and material

The datasets generated during the current study are available in the Pavlovia repository, https://gitlab.pavlovia.org/isonlab/hybrid_search

### Authors’ contributions

MJI and JEK conceived and designed the experiment. MJI produced the experiment code. AB collected the data. AB, MJI, and JEK performed the analyses. All authors discussed and interpreted the results. AB drafted the manuscript. All authors contributed to and approved the final manuscript.

## Notes

### Competing Interest Statement

The authors have declared no competing interest.

