## Supplemental Information for "Investigating the effects of contextual information, visual working memory, and inhibitory control in hybrid visual search"

*Change Detection Task (CDT) and Go-No Go (GNG) Procedure and Formulas*

**Figure S1.**

*CDT model trial of set size 4 with a change in the test array.*

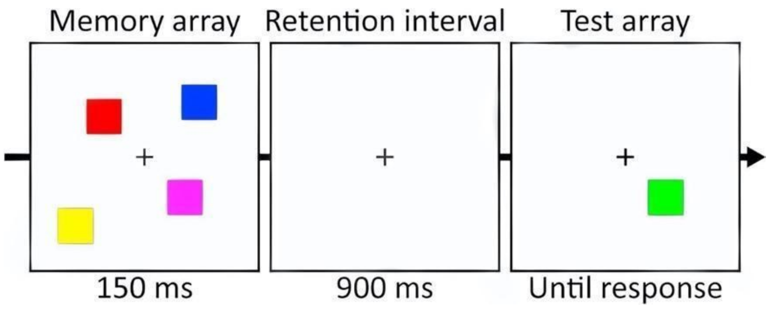


$$K = N \times (HR - FA)$$

$$HR = \frac{\text{\#correct responses in change trials}}{\text{\#change trials}}$$

$$FA = \frac{\text{\#incorrect responses in no-change trials}}{\text{\#no-change trials}}$$

Where N denotes the number of trials, HR is the hit rate and FA the false alarms rate.

**Figure S2.**

*GNGT example of trials throughout time.*

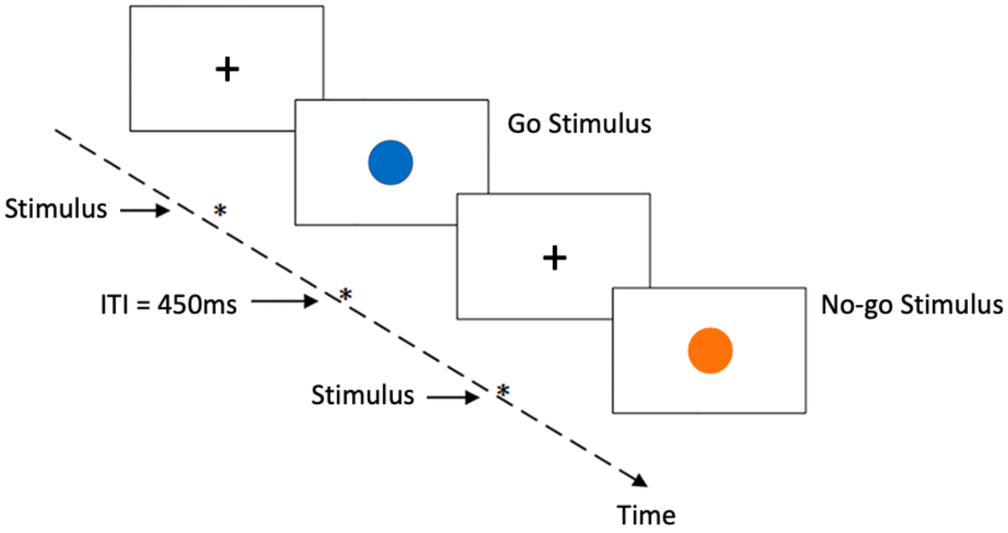

$$c = -\frac{1}{2}\left[z(HR) + z(FA)\right]$$

$$HR = \frac{\text{\#go response in go trials}}{\text{\#go trials}}$$

$$FA = \frac{\text{\# go response in no-go trials}}{\text{\# no-go trials}}$$

Here, c is the response bias. Z(HR) the z-transformed hit rate and Z(FA) the z-transformed false alarm rate. When FA are 0 or 1, then we adopted the corrections suggested by Stanislaw & Todorov, 1999 (substituting 0 for 0.5/N, and 1 for 1-0.5/N, where N is the number of target absent trials).

*Context Effects in Hybrid Search*

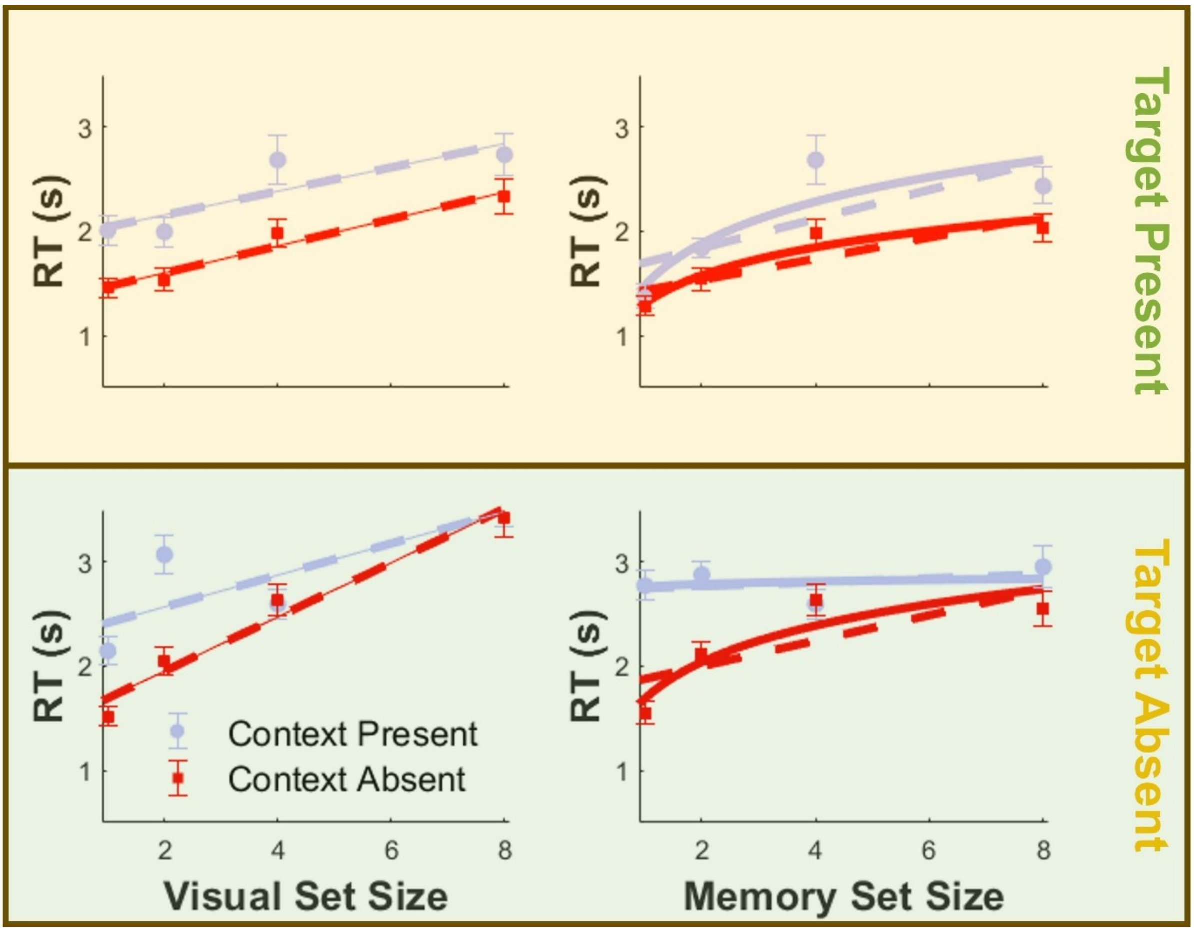


**Figure S3.** *RT as a function of visual and memory set sizes for TP, TA, CP and CA conditions for correct-only trials.* Error bars denote 95% CI. Dashed lines depict linear fits, while continuous lines portray logarithmic fits. Panels differentiate conditions: A) Visual Search Target Present; B) Visual Search Target Absent; C) Memory Search Target Present; D) Memory Search Target Absent.

**Table S1.**
Simple main effects analysis for VSS, target presence, and context present interaction on correct-trial RT

| VSS | Target Presence | Context Presence | Mean | F (1,71) | $\eta_p^2$ |
|---|---|---|---|---|---|
| 1 | TA | CA | 1.54 | 58.58** | .45 |
| | | CP | 2.11 | | |
| | TP | CA | 1.48 | 47.02** | .40 |
| | | CP | 1.99 | | |
| 2 | TA | CA | 1.99 | 94.35** | .57 |
| | | CP | 3.04 | | |
| | TP | CA | 1.53 | 19.48** | .22 |
| | | CP | 1.98 | | |
| 4 | TA | CA | 2.68 | 1.14 | .02 |
| | | CP | 2.58 | | |
| | TP | CA | 1.97 | 22.00** | .24 |
| | | CP | 2.66 | | |
| 8 | TA | CA | 3.43 | .485 | .01 |
| | | CP | 3.50 | | |
| | TP | CA | 2.40 | 6.84* | .09 |
| | | CP | 2.74 | | |

*Note.* TA = target absent; TP = target present; CA = context absent; CP = context present. Pillai's Trace is reported. P-values ≤.05 = *; p-values ≤ .001 = **.

**Table S2.**
Simple main effects analysis for MSS, target presence, and context present interaction on correct-trial RT

| MSS | Target Presence | Context Presence | Mean | F (1,74) | $\eta_p^2$ |
|---|---|---|---|---|---|
| 1 | TA | CA | 1.53 | 221.95** | .75 |
| | | CP | 2.76 | | |
| | TP | CA | 1.23 | 7.26* | .09 |
| | | CP | 1.37 | | |
| 2 | TA | CA | 2.12 | 111.47** | .60 |
| | | CP | 2.84 | | |
| | TP | CA | 1.54 | 13.37** | .15 |
| | | CP | 1.80 | | |
| 4 | TA | CA | 2.66 | .97 | .01 |
| | | CP | 2.56 | | |
| | TP | CA | 1.96 | 22.40** | .23 |
| | | CP | 2.64 | | |
| 8 | TA | CA | 2.56 | 10.24* | .12 |

| | | | | | |
|---|---|---|---|---|---|
| | | CP | 2.92 | | |
| | TP | CA | 2.02 | 18.16** | .20 |
| | | CP | 2.52 | | |

*Note.* TA = target absent; TP = target present; CA = context absent; CP = context present. Pillai's Trace is reported. P-values ≤ .05 = *; p-values ≤ .001 = **.

**Table S3.**
Simple main effects analysis for VSS, target presence, and context present interaction on accuracy.

| VSS | Target Presence | Context Presence | Mean | F (1,109) | $\eta_p^2$ |
|---|---|---|---|---|---|
| 1 | TA | CA | .85 | .78 | .01 |
| | | CP | .87 | | |
| | TP | CA | .73 | .19 | .00 |
| | | CP | .74 | | |
| 2 | TA | CA | .61 | 12.04** | .10 |
| | | CP | .70 | | |
| | TP | CA | .71 | 25.86** | .19 |
| | | CP | .55 | | |
| 4 | TA | CA | .72 | 21.33** | .16 |
| | | CP | .78 | | |
| | TP | CA | .67 | 4.77* | .04 |
| | | CP | .70 | | |
| 8 | TA | CA | .64 | 17.10** | .14 |
| | | CP | .53 | | |
| | TP | CA | .58 | 7.42* | .06 |
| | | CP | .49 | | |

*Note.* TA = target absent; TP = target present; CA = context absent; CP = context present. Pillai's Trace is reported. P-values ≤.05 = *; p-values ≤ .001 = **.

**Table S4.**
Simple main effects analysis for MSS, target presence, and context present interaction on accuracy

| MSS | Target Presence | Context Presence | Mean | F (1,109) | $\eta_p^2$ |
|---|---|---|---|---|---|
| 1 | TA | CA | .95 | 3.85 | .03 |
| | | CP | .91 | | |
| | TP | CA | .93 | .67 | .01 |
| | | CP | .92 | | |
| 2 | TA | CA | .92 | .10 | .00 |
| | | CP | .92 | | |
| | TP | CA | .59 | 13.45** | .11 |
| | | CP | .68 | | |
| 4 | TA | CA | .67 | 6.32* | .06 |
| | | CP | .70 | | |
| | TP | CA | .65 | 9.68* | .08 |
| | | CP | .60 | | |
| 8 | TA | CA | .44 | 20.35** | .16 |

| | | | | | |
|---|---|---|---|---|---|
| | | CP | .57 | | |
| | TP | CA | .58 | .02 | .00 |
| | | CP | .58 | | |

*Note.* TA = target absent; TP = target present; CA = context absent; CP = context present. Pillai's Trace is reported. P-values ≤.05 = *; p-values ≤ .001 = **.

### *Individual Differences in Hybrid Search*

**Table S5.**
*Simple Linear Regressions of K and c on accuracy and false alarm rates*

| **Linear Regressions** | **Std. Coeff. and t-values** | **$R^2$ and Model Sig.** |
|---|---|---|
| K → Accuracy | .12 (1.29) | .02 |
| K → Accuracy TP/CA | .10 (1.01) | .01 |
| **K → Accuracy TP/CP** | **.24 (2.55) *** | **.06*** |
| K → Accuracy TA/CA | -.06 (-.60) | .003 |
| K → Accuracy TA/CP | -.04 (-.39) | .001 |
| c → Accuracy | -.05 (-.48) | .002 |
| c → Accuracy TP/CA | .06 (.62) | .004 |
| c → Accuracy TP/CP | .01 (.14) | .00 |
| c → Accuracy TA/CA | -.09 (-.99) | .01 |
| c → Accuracy TA/CP | -.10 (-1.06) | .01 |
| c → FAs | .11 (1.15) | .01 |
| c → FAs TA/CA | .09 (.985) | .01 |
| c → FAs TA/CP | .10 (1.06) | .01 |

*Note.* Standardised β coefficients are the first values reported, followed by t-values inside brackets. P-values < .05 =*; p-values <.001 =**; Bold helps pinpoint significant models and IVs.
